# Trait diversity metrics can perform well with highly incomplete datasets

**DOI:** 10.1101/2022.11.08.515633

**Authors:** Kerry Stewart, Carlos P. Carmona, Chris Clements, Chris Venditti, Joseph A. Tobias, Manuela González-Suárez

## Abstract

1. Characterizing changes in trait diversity at large spatial scales provides insight into the impact of human activity on ecosystem structure and function. However, the approach is often based on trait datasets that are incomplete and unrepresentative, with uncertain impacts on trait diversity estimates.
2. To address this knowledge gap, we simulated random and biased removal of data from a near complete avian trait dataset (9579 species) and assessed whether trait diversity metrics were robust to data incompleteness with and without using imputation to fill data gaps. Specifically, we compared two commonly used metrics each calculated with two methods: trait richness (calculated with convex hulls and trait probabilities densities) and trait divergence (calculated with distance-based Rao and trait probability densities).
3. Without imputation, estimates of global avian trait diversity (richness and divergence) were robust when 30-70% of species had missing data for four out of 11 continuous traits, depending on severity of bias and the method used. However, when missing traits were imputed based on present morphological trait data and phylogeny, trait diversity metrics consistently remained representative of the true value, even when 70% of species were missing data for four out of 11 traits and data were not missing at random (biased with respect to body mass). Trait probability densities and distance-based Rao were particularly robust to missingness and bias when combined with imputation, with convex hull-based trait richness being less reliable.
4. Expanding global morphometric datasets to represent more taxa and traits, and to quantify intraspecific variation, remains a priority. In the meantime, our results show that widely used methods can successfully quantify large-scale trait diversity even when data are missing for two-thirds of species, so long as missing traits are estimated using imputation.

## Introduction

Trait-based approaches have emerged as an effective tool for addressing pressing questions in ecology and conservation (Green et al., 2022) and have been used to monitor ecosystem service delivery (Hevia et al., 2017), understand threats to ecosystem processes (Donoso et al., 2020), and forecast changes to ecosystem functioning (ŞekercioClu et al., 2004). In some cases assessments of trait diversity provide greater explanatory power than complementary measures of diversity such as species richness (Cadotte et al., 2011; Wilkes et al., 2020), in part due to the mechanistic link between traits and ecosystem functions (Petchey & Gaston, 2006). Assessing trait diversity can contribute advances necessary for tackling human-induced biodiversity loss (Mason & De Bello, 2013) through producing early warning systems of ecosystem collapse (Frainer et al., 2021, although see O’Brien et al., 2022 [in press]) and enhancing prioritisation of conservation efforts (Carmona et al., 2021; Cooke et al., 2019). Research into how trait diversity has varied across time and space could yield fruitful insights into ecosystem responses to global change (Green et al., 2022; Petchey & Gaston, 2006). Given the potential and versatility of trait-based approaches, ensuring trait diversity analyses are appropriately applied is of great importance (Blonder et al., 2018).

Traditionally, assessments of trait diversity have been carried out on local community-scales (Díaz & Cabido, 2001); only recently has the accumulation of large trait datasets enabled application on continental to global scales (Etard et al., 2020; Migliavacca et al., 2021). Large-scale trait analyses provide unique opportunities which complement those of local analyses. Integration of trait diversity across scales (Carmona et al., 2016), inclusion of species overlooked by local studies (such as migratory species) (Somveille et al., 2013), and identification of synergies and differences in the way humans impact natural ecosystems (Toussaint et al., 2021) provides a global perspective on biodiversity loss, needed for international policy development (Santini et al., 2021; Scholes & Biggs, 2005).

Large scale analyses come with their own set of challenges. Studying trait diversity at this scale requires relying on data that are rarely complete (González-Suárez et al., 2012; Toussaint et al., 2021), introducing uncertainty into inferences obtained (Richter et al., 2021). For example, while 80% of plant species are represented in the largest plant trait dataset (the TRY database) (Kattge et al., 2020), only 17% of species have data for 10 or more traits (Tobias et al., 2022), less than 1% had complete data across the six fundamental traits that were used to characterize the global spectrum of plant form and function (Díaz et al., 2016), and this percentage goes down to 0.1% when fine-root traits are also considered (Carmona et al., 2021). Etard et al. (2020) found that mean trait completeness across seven ecological traits was only 47% for amphibians and 46% for reptiles. Trait data availability is further restricted in invertebrates, fungi, algae, protozoa, and prokaryotes (Gallagher et al., 2019). In some groups the data availability is better. Etard et al. (2020) found that a mean of 89% of mammals and 84% of birds were represented (across seven ecological traits). A recent release of a new dataset, AVONET, sees 95-99% of bird species represented (with some variation by trait and taxonomy) for 11 morphological traits (Tobias et al., 2022) with ecological and geographical data also available for most species. However, even when a small proportion of species are missing data, the available information may not be representative of the true diversity (González-Suárez et al., 2012). For most taxonomic groups, data are not missing at random, and are biased according to morphological, ecological, and geographical traits (González-Suárez et al., 2012; Sandel et al., 2011; Tyler et al., 2012). This may affect the accuracy of trait diversity metrics calculated (Sandel et al., 2011), potentially leading to spurious or misleading results (Pincheira-Donoso & Hodgson, 2018) and impeding the application of trait-based methods (Etard et al., 2020).

Emerging methods to estimate trait diversity can now be applied large-scale trait databases (Blonder et al., 2018; Carmona et al., 2016) but there is uncertainty over how these methods respond to missing data, particularly when data are not missing at random (Petchey & Gaston, 2006). This lack of understanding risks making unfounded assumptions that data are representative (Tyler et al., 2012) and impedes correct application of trait diversity methods (Pakeman et al., 2014). Where continuous trait data are available, functional diversity is most commonly assessed using three broad methods: distance-based methods, convex hulls, and probabilistic hypervolumes (Mammola et al., 2021). Using these methods distinct aspects of functional diversity can be described including richness (the total volume of trait space occupied by an assemblage) and divergence (the variance in traits of biological units in an assemblage) (Mason et al., 2005). A few studies have quantified sensitivity to missing data at local scales using distance-based methods (Pakeman, 2014; Richter et al., 2021). However, to our knowledge no one has quantitatively compared methods spanning conceptual frameworks (Mammola et al., 2021). This is urgently needed because local scale sensitivity and bias may differ substantially to those that occur at large scales and sensitivity may differ when using multidimensional methods.

The best approach for dealing with missing data is debated (Taugourdeau et al., 2014). One approach is to remove all species with missing data, known as complete case analysis (Freckleton, 2008). Alternatively, missing data can be imputed. While in some instances complete case analysis can outperform imputation (Johnson et al., 2021), research has also documented the danger of introducing bias into the results, as data are often not missing at random (Freckleton, 2008). The performance of different imputation methods has been evaluated in several recent studies (Johnson et al., 2021; Penone et al., 2014) that collectively identify most reliable methods, and for trait datasets, agree on the value of using phylogenetic information to improve accuracy (Debastiani et al., 2021). However, whether imputation is needed or valuable when estimating trait diversity remains largely unexplored. Here, we test how data missingness and bias affect estimation of trait diversity metrics using both complete-case analysis and imputation. In particular, we address three goals: 1) quantify changes due to missing and biased data in two types of diversity metrics: trait richness and trait divergence, each estimated with two methods; 2) determine if data imputation methods reduce errors and biases in estimated metrics, and 3) identify the degree of missingness and bias permissible when quantifying large-scale trait diversity. In contrast with many previous studies that use simulated datasets, here we take advantage of the recently released near-complete bird trait dataset, AVONET (Tobias et al., 2022). Using a real dataset offers a more realistic representation of empirical diversity analyses, where outliers and non-normal distributions are likely to be present (Junker et al., 2016). Our findings assist methodological decision-making in trait diversity analyses providing direct recommendations on how to proceed when using incomplete trait datasets.

## Methods

### The master dataset

Using the AVONET dataset of avian morphological traits (Tobias et al., 2022), we investigated the impact of data missingness and bias in estimates of trait richness and trait divergence. In this analysis, we followed the BirdTree taxonomy and phylogeny (Jetz et al., 2012). From the 9993 species recognized in BirdTree, empirical data are missing for one or more traits for 410 species which were removed from the analysis. As this is <5% of species this is not expected to have a considerable impact on our findings. All morphological traits were included: beak length (culmen), beak length (nares), beak width, beak depth, tarsus length, wing length, secondary length, tail length, body mass, hand wing index, and Kipps distance (for additional details on how traits were measured see, Tobias et al., 2022). All trait values were log_10_ transformed and standardised. While we aimed to use all available data, after initial exploration we removed the four species of kiwis (Apterygidae) recognised in the BirdTree phylogeny (Jetz et al., 2012) as they were extreme outliers with respect to morphological traits (see Supplementary Information section 1, Matthews et al., 2022). The final master dataset included data for 9579 bird species.

### Overview

From the master dataset we created 2800 datasets by removing data under all permutations of seven degrees of missingness, four degrees of bias and two ways of dealing with missing data (complete case analysis and imputation), with 50 replicates for each permutation (for each replication, data were removed at random under stipulated missingness and bias conditions). Principal component analysis was used for dimensionality reduction. For every dataset we calculated four metrics: convex hull (trait richness), distanced-based Rao (trait divergence), and trait probability densities (trait richness and divergence). Figure 1 provides a schematic overview of the methodology.

**Figure 1.**
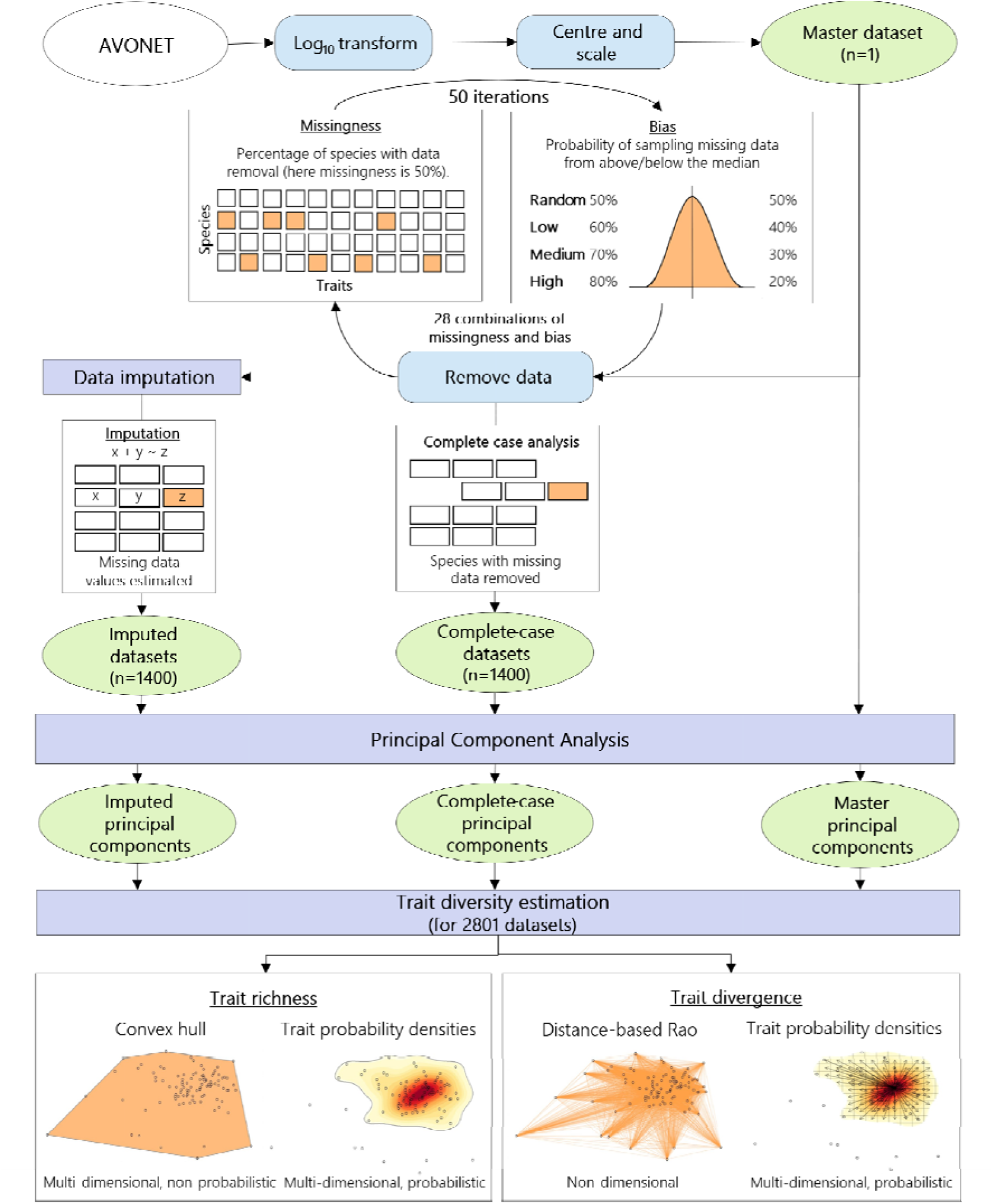
Summary of the method of this analysis.

### Missingness

Missingness refers to the percentage of species from which data were removed. We applied seven degrees of missingness, from 10% to 70%, at 10% intervals. The main text shows results based on removing data from four out of 11 traits for each species with data removal (chosen at random). Supplementary Information section 2 shows results based on removing values from seven out of 11 traits.

### Bias

We emulated a commonly observed bias in trait data missingness, wherein species with lower body masses are more likely to be missing data (Tyler et al., 2011; González-Suárez et al., 2012; Garamszegi and Møller; 2013). Bias was applied by increasing the probability that data were removed from species with a below median body mass with four scenarios: no bias – 50% probability, low bias – 60% probability, medium bias – 70%, high bias – 80%. This means that in a low bias scenario, there was 60% probability that data were removed from species with below median body mass, and 40% probability that data were removed from species with above median body mass. The extent of bias in these scenarios was chosen based on analysis of bias in historic accumulation of data in large-scale bird trait datasets (see Supplementary Information section 3).

### Imputation

We tested two ways of dealing with the generated incomplete datasets: 1) removal of species with missing data (complete case analysis), and 2) filling data gaps through imputation. We used missForest imputation, implemented through the *missForest* package (Stekhoven & Bühlmann, 2012), due to its demonstrated accuracy (Penone et al., 2014; Hong & Lynn, 2020), and fast computation times. Phylogenetic data were incorporated through eigenvectors (Debastiani et al., 2021). Evolutionary relationships were based on a maximum clade credibility tree from the first 1000 trees provided by Jetz et al. (2012). As biased removal of data may affect tree imbalance (Fischer et al., 2021) we tested the impact of biased data removal on tree balance and the impact of tree balance on imputation accuracy (Supplementary information section 4).

### Principal component analysis

For each dataset we carried out principal component analysis. Variation in bird morphology can be meaningfully explained in a small number of dimensions (Pigot et al., 2020) so we used only the first three principal components that collectively explained 90% of total variance for the master dataset.

Previous research has analysed the impact of dataset incompleteness on principal component analysis (Dray and Josse, 2014). Here we found that carrying out principal component analysis on incomplete datasets did not introduce a substantial component of error (Supplementary Information section 4).

### Trait diversity estimation

For each of the 2801 datasets we calculated four metrics, two estimating trait richness and two estimating trait divergence (figure 1, Table 1). Trait richness is a measure of the volume of trait space occupied (Mammola et al., 2021). We calculated trait richness using convex hull (a multidimensional, non-probabilistic method) and trait probability densities (a multidimensional probabilistic method). Trait divergence is a measure of the dispersion of trait space occupation (how far species are from each other, or the centre of trait space [Mouchet et al., 2010]). We calculated trait divergence using Rao (a nondimensional distance-based method), and trait probability densities. Distance-based methods (Rao) and convex hull are the methods underlying the widely used *FD* package (Laliberté & Legendre, 2010). We compared distance-based methods and convex hull to a more recent approach using probabilistic hypervolumes, implemented through trait probability densities in the package *TPD* (Carmona et al., 2016). Where user-defined parameters were available (alpha value in TPD) default parameterisations were used. We calculated trait diversity of the bird assemblage as a whole, using presence only data (no information on abundance). We did not consider intra-specific trait diversity or calculate overlap between taxonomic or ecological groups.

**Table 1.** Trait diversity methods used for quantifying functional diversity.

| METHOD | METRIC | DESCRIPTION | R FUNCTION<br>USED (FOR<br>BUILDING TRAIT<br>SPACE) | R FUNCTIONS USED<br>(FOR<br>CALCULATING<br>TRAIT DIVERSITY) |
| --- | --- | --- | --- | --- |
| **DISTANCE-**<br>**BASED**<br>**METRICS** | Trait<br>divergence | Non-dimensional | n.a. | melodic (de Bello et al.,<br>2016) |
| **CONVEX**<br>**HULL** | Trait richness | Binary hypervolume<br>(multidimensional) | BAT::hull.build<br>(Cardoso et al.,<br>2015) | BAT:: hull.alpha<br>(richness) (Cardoso et<br>al., 2015) |
| **TRAIT**<br>**PROBABILITY**<br>**DENSITIES**<br>**(TPD)** | Trait richness<br>and divergence | Probabilistic<br>hypervolume<br>(multidimensional) | TPD:: TPDs<br>(Carmona et al.,<br>2016) | TPD:: REND (richness,<br>evenness and<br>divergence) (Carmona et<br>al., 2016) |

### Statistical analysis

To determine the impact of missingness, bias and how missing data were handled (complete case analysis or imputation) we calculated the deviation in trait diversity metrics for each dataset replicate *i* for each trait method *m* as,

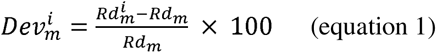

Where *Rd_m_* is trait diversity of the master dataset estimated using metric *m* and 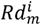 is the trait diversity of replicate dataset *i* estimated using the same metric *m*. Depending on the metric, trait diversity represents trait richness or trait divergence (figure 1).

We then fitted a multiple linear regression to explain *Dev* values as a function of missingness (continuous variable), bias (ordered factor), missing data handling (binary variable with values: complete-case and imputation) and the trait diversity method (categorical variable). The model was fitted using the *lm* function in base R version 4.1.2 (R Core Team, 2021) with all predictor combinations and their interactions. We evaluated the model using Analysis of Variance (ANOVA) (*anova* function in library *ape* [Paradis, Claude & Strimmer, 2004]). We define permissible levels of missingness as those where the deviation in estimated trait diversity (*Dev*) was <10% for all dataset replicates.

Additionally, we calculated imputation accuracy using normalised root mean square error (NRMSE); the mean of imputation errors divided by the trait range in the master dataset (equation 2). We calculated the mean NRMSE across all traits for every imputed dataset (n=1400).

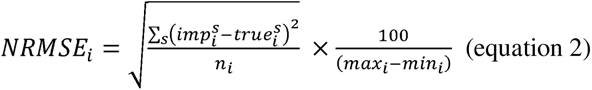

Where *NRMSE_i_* is the normalised root mean square error for trait *i*, 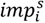 is the imputed value for species *s* in trait *i*, 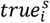 is the true value in the master dataset for species *s* in trait *i*, *n_i_* is the number of imputed values for trait *i* across all species and *max_i_* and *min_i_* are the maximum and minimum value for trait *i* in the master dataset, respectively.

## Results

Trait diversity estimates from incomplete datasets deviated from true values (calculated with the master dataset) by between −48% and 31%. Deviations were affected by all tested predictors and their interactions (table 2, R^2^=0.82). Significant interactions among predictors demonstrate the complexity of response to incomplete data, but by and large, trait diversity estimates deviated more from the true values at greater levels of missingness and bias (figure 2). Deviations were usually greater and more clearly affected by bias when only species with no missing data were analysed (complete case analysis) highlighting the benefit of imputation (figure 2). In the case of distance-based Rao, the absolute error with imputation was on average 5.2 times smaller than in the complete case scenario, and for convex hull the absolute error with imputation was on average 1.9 times smaller than then complete case scenario.

**Figure 2.**
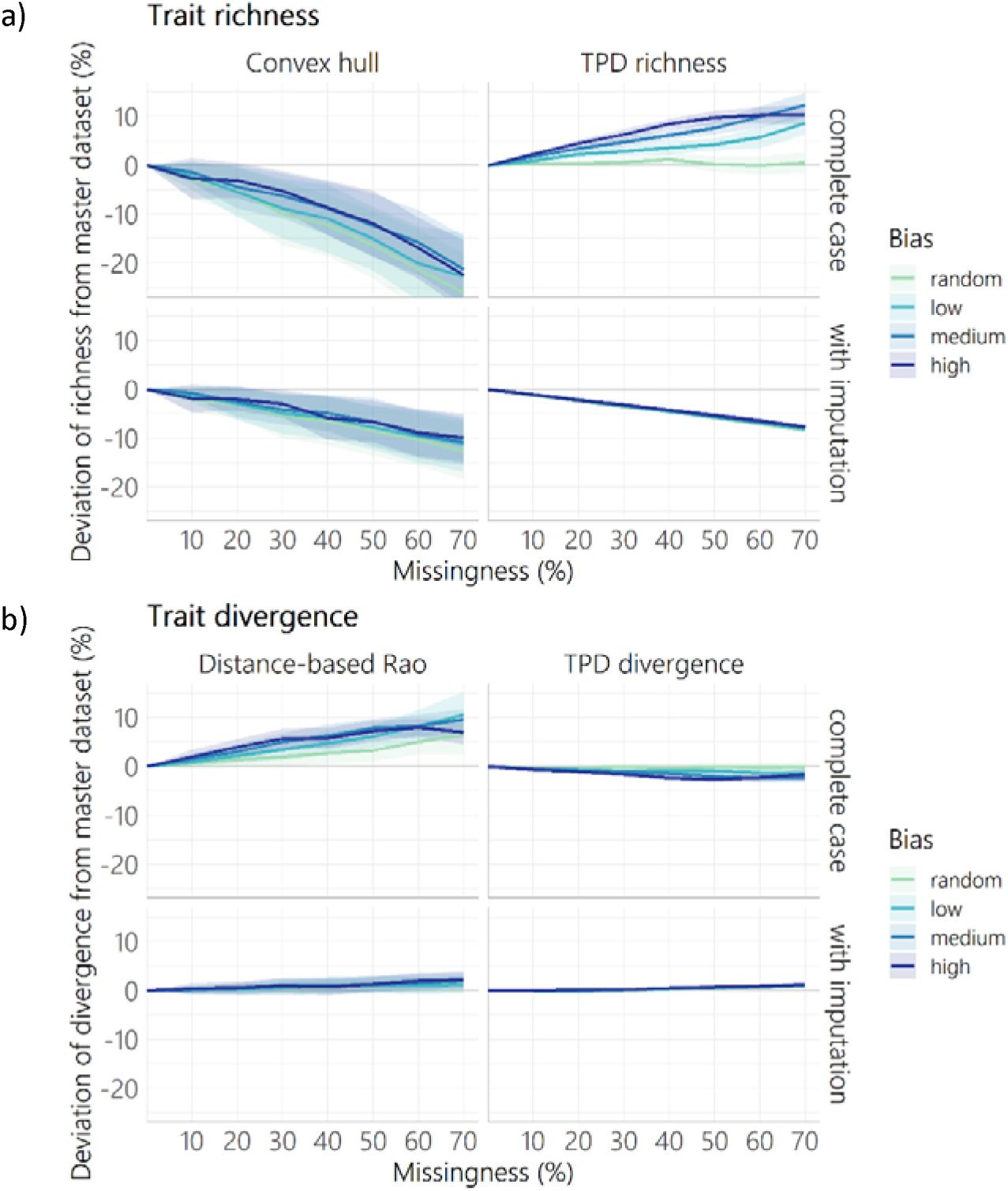
Deviation in a) trait richness and b) trait divergence for incomplete datasets compared to the master dataset given varying levels of missingness (percentage of species with missing data for four out of 11 traits) and bias (greater probability of data missing from small-sized species). Lines represent the mean with shaded areas showing 1 SD from the 50 replicates analysed for each combination. Top panels (in a and b) show results for complete case analyses (only species with all trait data analysed), and bottom panels show results for when missing values were imputed using random forest implemented with the missForest R function. Trait diversity was calculated using four metrics: convex hull (richness), trait probability densities (TPD) richness, distance-based Rao (divergence) and trait probability densities (TPD) divergence.

**Table 2.** Analysis of Variance (ANOVA) table of a linear model explaining Deviation (Dev) based on missingness, bias, trait diversity metric and way of handling missing data (and their interactions). Rows are ordered by the F value, with the strongest predictor (Metric) at the top. All predictors explained a significant proportion of variance (p value <0.001 for all predictors).

| Predictor | Df | Sum Sq | Mean Sq | F value | p value |
| --- | --- | --- | --- | --- | --- |
| **Metric** | 3 | 186447.0 | 62149.0 | 8525.2 | <0.001 |
| **Metric: Missingness** | 3 | 94994.2 | 31664.7 | 4343.6 | <0.001 |
| **Imputation: Metric** | 3 | 74670.6 | 24890.2 | 3414.3 | <0.001 |
| **Missingness** | 1 | 12690.9 | 12690.9 | 1740.9 | <0.001 |
| **Imputation: Metric: Missingness** | 3 | 35348.0 | 11782.7 | 1616.3 | <0.001 |
| **Imputation** | 1 | 5787.8 | 5787.8 | 793.9 | <0.001 |
| **Bias** | 3 | 4693.9 | 1564.6 | 214.6 | <0.001 |
| **Imputation: Missingness** | 1 | 1357.2 | 1357.2 | 186.2 | <0.001 |
| **Imputation: Bias** | 3 | 1985.0 | 661.7 | 90.8 | <0.001 |
| **Missingness: Bias** | 3 | 1335.8 | 445.3 | 61.1 | <0.001 |
| **Metric: Bias** | 9 | 3882.3 | 431.4 | 59.2 | <0.001 |
| **Imputation: Metric: Bias** | 9 | 3261.4 | 362.4 | 49.7 | <0.001 |
| **Imputation: Missingness: Bias** | 3 | 658.9 | 219.6 | 30.1 | <0.001 |
| **Metric: Missingness: Bias** | 9 | 1571.0 | 174.6 | 23.9 | <0.001 |
| **Imputation: Metric: Missingness: Bias** | 9 | 1440.2 | 160.0 | 22.0 | <0.001 |
| **Residuals** | 12736 | 92845.8 | 7.3 | NA | NA |

As an exception, the lowest deviations when using TPD richness occurred when data were missing at random and complete case analysis was used. TPD divergence never deviated from the true values by more than 5% regardless of bias, missingness, or the way of handling missing data (imputation or complete case analysis). Consistency within replicates (for the same level of missingness and bias) was variable among methods. Deviation was less predictable (more variation among replicates) when using complete case analyses, and when using convex hull to estimate trait richness.

Whether missing data resulted in over- or under-estimation of trait diversity was variable, and dependant on the method used and how missing data were handled. When species with missing data were removed, trait richness was underestimated with convex hull but overestimated with TPD. When using imputation, trait richness was underestimated with both convex hull and TPD. Distance-based methods consistently overestimated trait divergence when species with missing data were removed, whereas TPD divergence was slightly underestimated trait divergence. With imputation, the effect of missing data on estimates of trait divergence was negligible for both distance-based methods and TPD (figure 2).

Despite these varying patterns, if imputation was used, trait divergence from distance-based Rao was never overestimated by more than 6% and trait divergence from TPD was never overestimated by more than 3% (versus 31% overestimation and 5% underestimation in the complete case scenario, respectively). With imputation, trait richness from TPD was never underestimated by more than 10% (versus 19% overestimation in the complete case scenario). Trait richness from convex hull was underestimated by up to 48% without imputation and 24% with imputation. Metrics estimated using distance-based and TPD methods were insensitive to bias when imputation was used. When using complete case analysis, the extent of missing data permissible was dependant on the bias scenario and method, with biased data removal resulting in lower missingness permissible for all metrics other than TPD divergence. Trait divergence from TPD was extremely robust to missing and biased data and did not deviate by more than 10% from true values even at 70% missingness, regardless of whether complete case analysis or imputation was used. Convex hull was much less reliable with very limited levels of missing data permissible (figure 3). Lower missingness was permissible when seven out of 11 (rather than four out of 11) traits were removed except for trait divergence from TPD (see Supplementary Material section 2). Overall, these results indicate that when using imputation, TPD richness, TPD divergence and distance-based Rao (divergence) can be robustly estimated even with high data missingness (up to 70%) (given high trait coverage).

**Figure 3.**
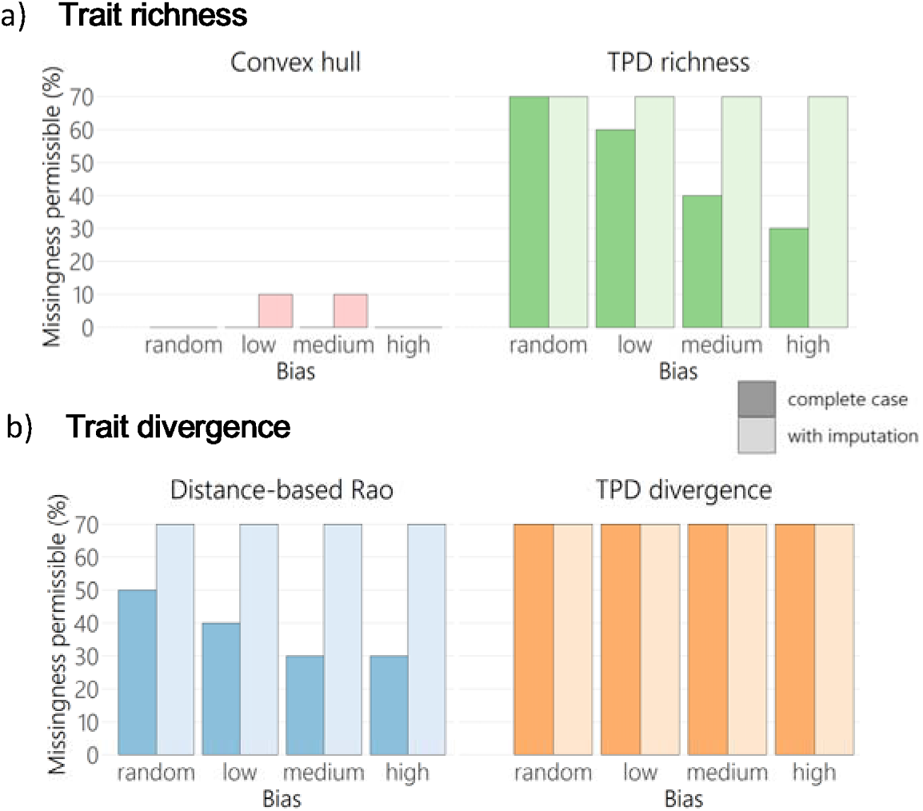
Maximum missingness permissible (maximum missingness where all trait diversity values fell within +/− 10% of trait diversity of the master dataset) for a) trait richness and b) trait divergence, with four degrees of bias. Trait diversity was calculated using convex hull (richness), trait probability densities (TPD) richness, distance-based Rao (divergence) and trait probability densities (TPD) divergence. Results are shown for complete case analysis (darker colour bars) and for imputed datasets (lighter colour bars).

Imputation was very accurate even when data removal was extreme and highly biased (imputation error was always <2.5% of the range, figure 4). Imputation error was lower with more bias, contrary to what we may have assumed, but not surprising given the distribution of data in the empirical dataset. Small bird species had higher clustering of trait values than large species (there are more small birds with more similar traits), facilitating more accurate imputation. Because in high bias replicates, values were more often removed from small species, this resulted in higher accuracy of imputation overall.

**Figure 4.**
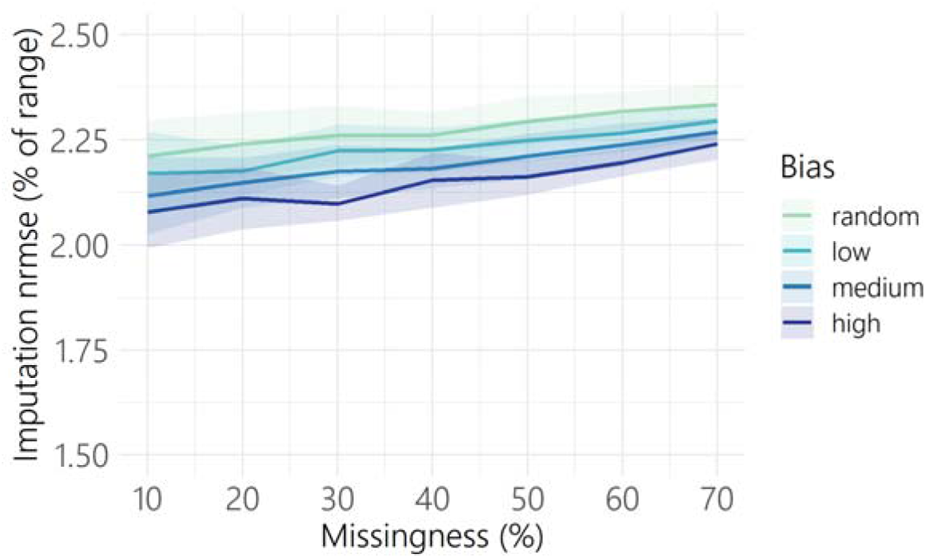
Imputation normalised root mean square error (expressed as a percentage of trait range, mean shown by line with shaded areas showing 1 SD from 50 replicates) with varying levels of missingness (percentage of species with missing data for four out of 11 traits) and bias (greater probability of data missing from small-sized species).

## Discussion

Our results indicate that basic metrics of trait richness and divergence can be relatively accurately estimated with incomplete data. From the methods we tested, we recommend estimating trait richness with TPD, although both TPD and distance-based Rao were reliable for trait divergence. We also recommend using imputation as we found that imputing missing trait data reduced deviations in trait diversity metrics to <10%. This finding is promising for the application of large-scale trait analyses. In global analyses, complete trait information is rare, especially as the number of traits used increases to provide a more comprehensive assessment of functional diversity. As such many studies rely on imputation, but the impact of this decision is often questioned. Our findings show that imputation can be an effective way for accounting for incomplete data, and in our avian dataset greatly reduced errors in functional diversity estimation compared to removing species with incomplete data. While differences in the number of traits, and application of trait diversity metrics must be considered, our results suggest that given observed levels of missing data for many taxa (Etard et al., 2020, Kattge et al., 2020, Tobias et al., 2022) current methods could be used to calculate trait diversity with reasonable accuracy.

Despite the robustness of trait diversity metrics (aside from trait richness estimated using convex hull), careful consideration is needed when interpreting trait diversity values, as some unexpected responses to incomplete data were observed. Trait richness is the volume of trait space occupied, so it is expected to shrink with data removal. While this was observed for convex hull, TPD richness increased with biased removal of species when using complete case analysis. This counterintuitive response is due to data-defined bandwidth selection during kernel density estimation of trait probability densities (see Supplementary Information section 6 for detailed explanation). These results emphasise the need for null models and clear consideration of expectations when conducting trait diversity analyses (particularly when estimating trait richness). Outlining expectations and using null models ensures results are interpreted correctly and observed responses are due to the hypothesis being tested rather than an artefact of data availability or metric behaviour.

Our comparison of trait diversity methods revealed sensitivities to missing data that were previously unreported. Convex hulls performed consistently worse than TPD richness, with low tolerance to missing data even when imputation was used. This suggests that when relying on incomplete datasets to quantify trait richness, using TPD (or similar approaches based on probabilistic hypervolumes [Blonder, 2017]) is advisable. This does not preclude value in convex hull approaches. Convex hulls are sensitive to changes which affect extreme trait values, so if a complete dataset is available, they can provide an estimation of trait diversity which is highly responsive to changes occurring in an ecosystem. Our findings agree with others that sensitivity to missingness and bias varies among methods (Pakeman, 2014) and attest to the robustness of distance-based metrics such as Rao (Májeková et al., 2016; Richter et al., 2021). We also found that both trait divergence metrics; distance-based Rao and TPD, were particularly robust to incomplete data. TPD (divergence and richness) and distance-based methods, if used with accurate imputation, can be applied to large-scale trait datasets with confidence, even at moderate to high levels of missingness and bias.

For all methods, we found that imputation was generally effective for overcoming gaps and biases in incomplete datasets, even at extreme levels of missingness and bias. This is in agreement with previous studies (Johnson et al., 2021; Penone et al., 2014; Swenson, 2014) and highlights the value of imputation when appropriately applied. In our analysis, imputation was particularly effective at eradicating the signal of bias. This is important given that bias in missing data are prevalent in large-scale datasets (González-Suárez et al., 2012; Sandel et al., 2011; Tyler et al., 2012) and the extent of bias is often unknown (Tyler et al., 2012). Despite this, in some cases, imputation could lead to greater errors than complete case analysis (see also Johnson et al., 2021). Therefore, while our results agree with others that random forest models (as implemented by the *missForest* R function) are an accurate imputation method for trait data (Penone et al., 2014; Hong & Lynn, 2020), care should be taken to ensure use of imputation is appropriate. Our findings regarding the utility of imputation are only applicable to continuous trait imputation, as the efficacy of categorical traits imputation was not explored. In our study, even under high missingness, imputation was based on a large number of species with a well-defined and complete phylogeny, and data was missing for only four out of 11 traits (or seven out 11 traits, see Supplementary material section 4). The utility of imputation in tackling missing and biased data has been shown to depend on the correlation between traits, and extent of phylogenetic autocorrelation (Clavel et al., 2014). As there is a strong link between form and function in birds (Pigot et al., 2020) phylogenetically informed imputation methods are expected to be particularly reliable for this group. Studies should assess imputation accuracy, through methods such as bootstrapping (Khan et al., 2019) and testing the ability to locate species within trait space (Carmona et al., 2020), to ensure imputation is adequately accurate for the desired application. In addition, when data are only available for a small subset of taxa or traits, or phylogenies are unavailable, we recommend reducing the scope of the study to focus on a smaller group with better data availability.

Making an informed decision on how much missing data can be tolerated is one of the challenges of trait diversity analyses. When using imputation and TPD or distance-based methods, we found up to 70% missingness (the maximum missingness simulated) was permissible under our criterion of trait diversity estimates falling within 10% of true values. However, the maximum missingness permissible was lower when using complete case analysis and was very low with convex hull trait richness. For smaller datasets, when using less-well-defined phylogenies or when many dimensions are needed to quantify trait space (principal components in this case), the maximum missingness permissible will likely be reduced. When using TPD richness and imputation, and removing seven out of 11 traits values rather than four, the maximum missingness permissible (where deviation of richness from true values was less than 10%) was reduced to 40% (Supplementary Material section 6). Therefore while 70% missingness can serve as a guideline where application is similar to that reported here, the spatial scale of the analysis, the size of the taxonomic or ecological group, the correlation between traits, and the extent of phylogenetic conservatism could affect maximum missingness permissible in other contexts. In addition, more restricted criteria for acceptable deviation from true values (defined as 10% here) may be desirable in some cases leading to lower missingness permissible. Future work is needed to explore how different factors influence trait diversity estimation when using incomplete datasets. In the meantime, wherever possible the extent of missingness should be quantified and used alongside other factors to inform choice of imputation and trait diversity metric.

Trait-based analyses are increasingly being applied to greater taxonomic and spatial scales (Etard et al., 2020, Johnson et al., 2011), so it is likely that trait-based ecologists will have to deal with missing data for the foreseeable future. Our findings show that imputation can permit reasonably accurate measurements of functional diversity even when 70% of species have incomplete data. In particular, TPD methods (for richness and divergence metrics) and distance-based methods (Rao for divergence) resulted in robust quantification of trait diversity. Trait richness estimated using convex hull was sensitive to incomplete data and should be used with caution. Given observed deviations we can conclude that for some taxa and traits, we have enough data to perform accurate large-scale analyses of trait diversity. Overall, the findings presented here are promising for the application of functional diversity methods to large-scale questions in ecology and conservation, and highlight the potential of imputation for overcoming gaps in trait databases.

## Data availability

AVONET (Tobias et al., 2022) is available for use under the creative commons licence (CC BY 4.0) and can be accessed at: https://figshare.com/s/b990722d72a26b5bfead.

BirdTree phylogenetic and taxonomic data can accessed at: https://birdtree.org/.

Simulated incomplete datasets and data underlying figures and tables will be made available on the University of Reading research data archive (https://researchdata.reading.ac.uk/).

## Acknowledgements

KS acknowledges PhD studentship funding from the SCENARIO NERC Doctoral Training Partnership grant NE/S007261/1. CPC was supported by the Estonian Ministry of Education and Research (PSG293) and the Mobilitas Pluss programme of the Estonian Research Council (MOBERC40). CC is supported by grant NE/T006579/1. CV is supported by the Leverhulme Trust (RL-2019-012).

## Author contributions

All authors conceived the ideas and designed methodology; JAT compiled the data; KS analysed the data with substantial contributions from CPC; KS led the writing of the manuscript with substantial contributions by MGS. All authors contributed critically to the drafts and gave final approval for publication.

## Supplementary Information

### 1. The kiwi problem: convex hull sensitivity to outliers

When this analysis was first carried all four extant kiwi species (Apterygidae) were included. The distribution of species along the first two principal components shows that kiwis are outliers in this trait space (figure S1), which greatly impacted the trait richness estimates based on convex hulls (figure S2).

**Figure S1.**
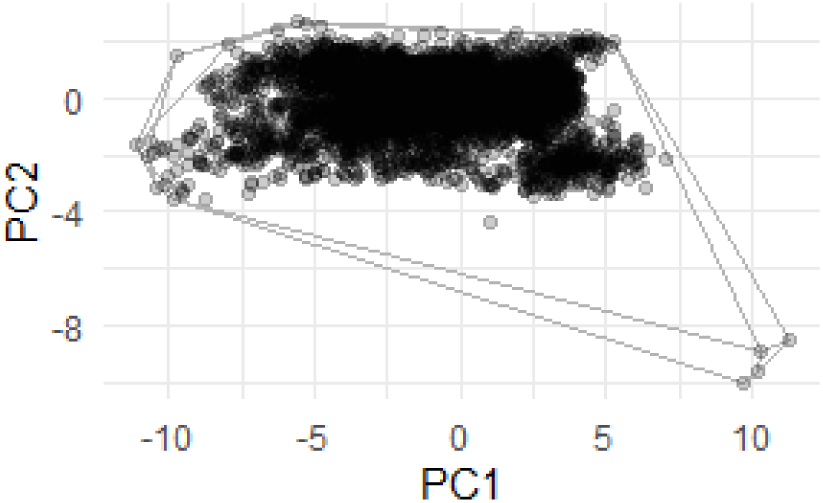
Distribution of bird species along the first two principal components (PC1 and PC2) (points) showing convex hull (hull). This is for a scenario where 10% of species were removed at random (r10). The four points on the bottom right are the four extant kiwi species.

The presence of outliers meant that trait richness (when calculated using convex hulls) responded in unexpected and strong ways to data removal, particularly when data was removed at random (figure S2).

**Figure S2.**
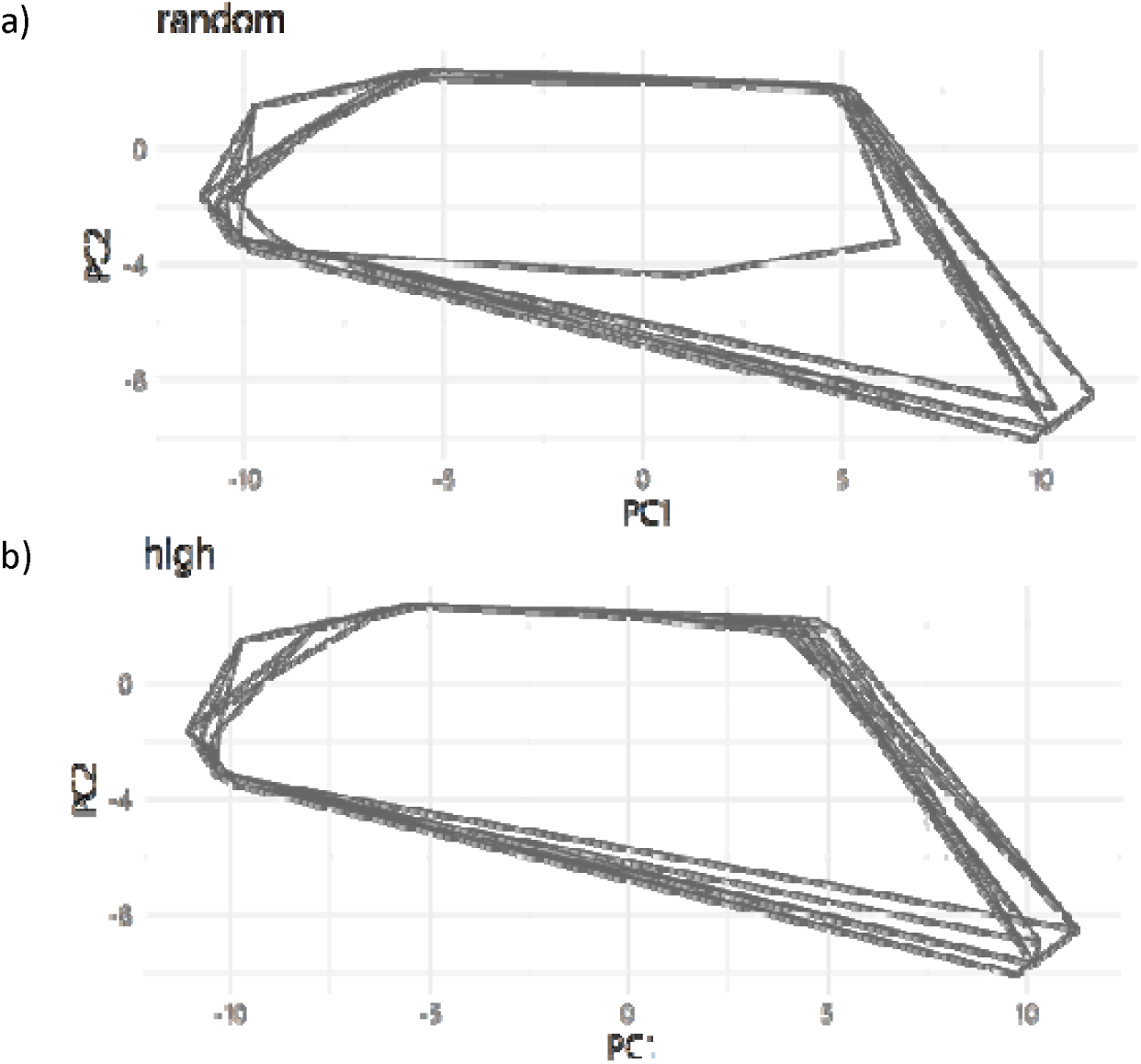
Convex hulls of bird species with 50% missingness, for an a) random and b) high bias scenario. 10 replicates for each scenario are shown.

When data were removed at random, the probability of all kiwi species being removed was greater than when data removal was biased, as the kiwi species have above median body masses. Where all kiwis were removed there was a trait richness decreased substantially giving high deviation from the master dataset, with higher deviations for random than biased data removal. Because of this we chose to discard kiwis and rerun the analysis.

These findings highlight the sensitivity of convex hull to outliers. Without use of a real dataset, it is unlikely that this sensitivity would have been revealed.

### 2. Removing seven out of 11 traits

We repeated the analyses removing seven out of 11 traits for each species with data removal. Deviation from the master dataset was greater than when four out of 11 traits were removed for all metrics apart from TPD divergence which remained robust to incomplete data. Despite greater deviation for most metrics, differences between methods and the effect of using imputation remained the same (figure S3).

**Figure S3.**
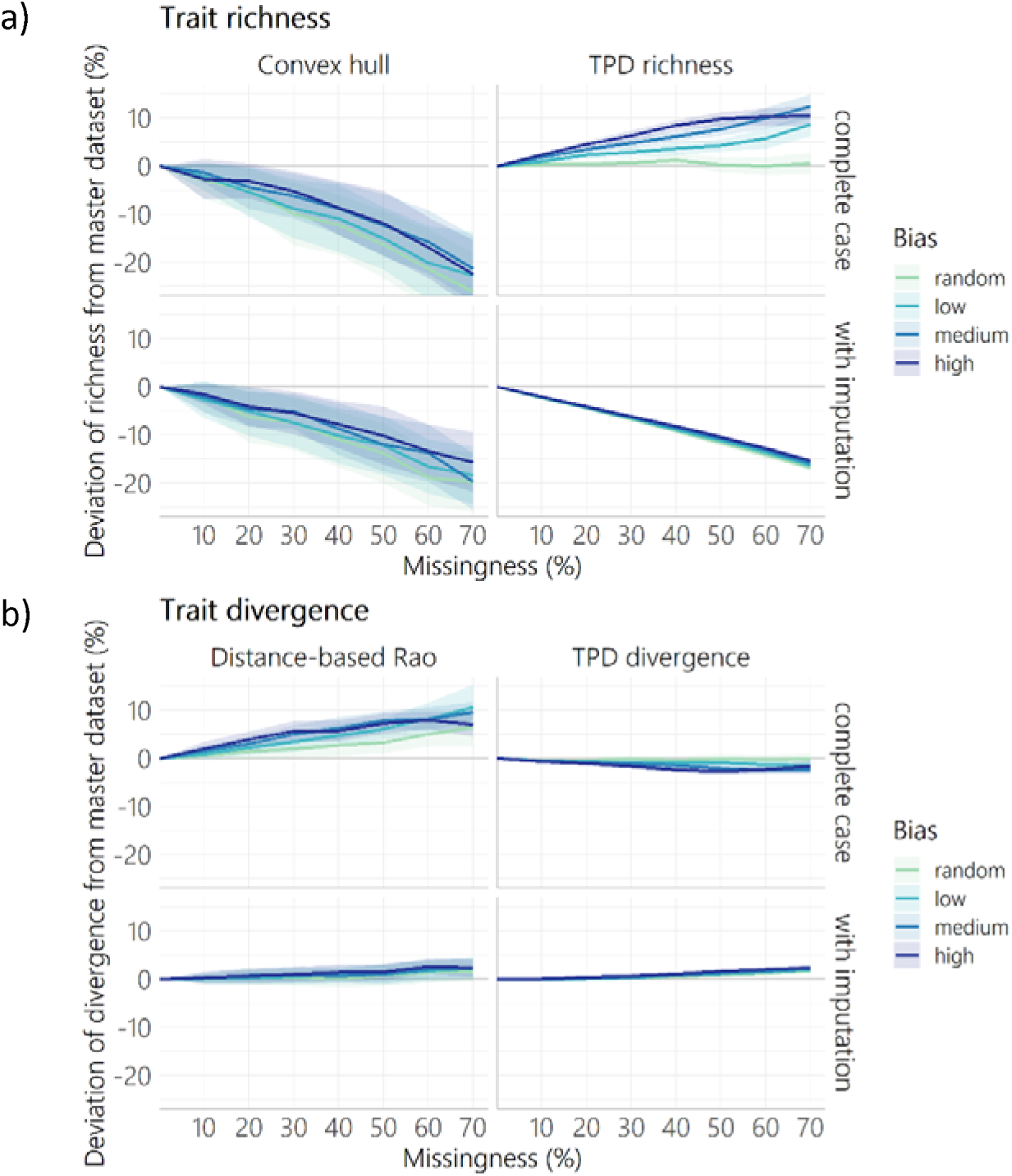
Deviation in a) trait richness and b) trait divergence for incomplete datasets compared to the master dataset given varying levels of missingness (percentage of species with missing data) and bias (greater probability of data missing from small-sized species). For each species with missing data seven out of 11 trait values were removed at random. Lines represent the mean with shaded areas showing 1 SD from the 50 replicates analysed for each combination. Top panels (in a and b) show results for complete case analyses (only species with all trait data analysed), and bottom panels show results for when missing values were imputed using random forest implemented with the missForest R function. Trait diversity was calculated using four metrics: convex hull (richness), trait probability densities (TPD) richness, distance-based Rao (divergence) and trait probability densities (TPD) divergence.

Removing seven out of 11 traits reduced missingness permissible when using TPD richness with imputation (30-40% missingness permissible compared to 70% when only four out of 11 traits were removed, figure S4). Trait divergence metrics (distance-based Rao and TPD divergence) were more robust to less trait information, and missingness permissible with imputation remained at 70%, even in high bias scenarios. No missing data (lowest tested value was 10%) was permissible when using convex hull.

**Figure S4.**
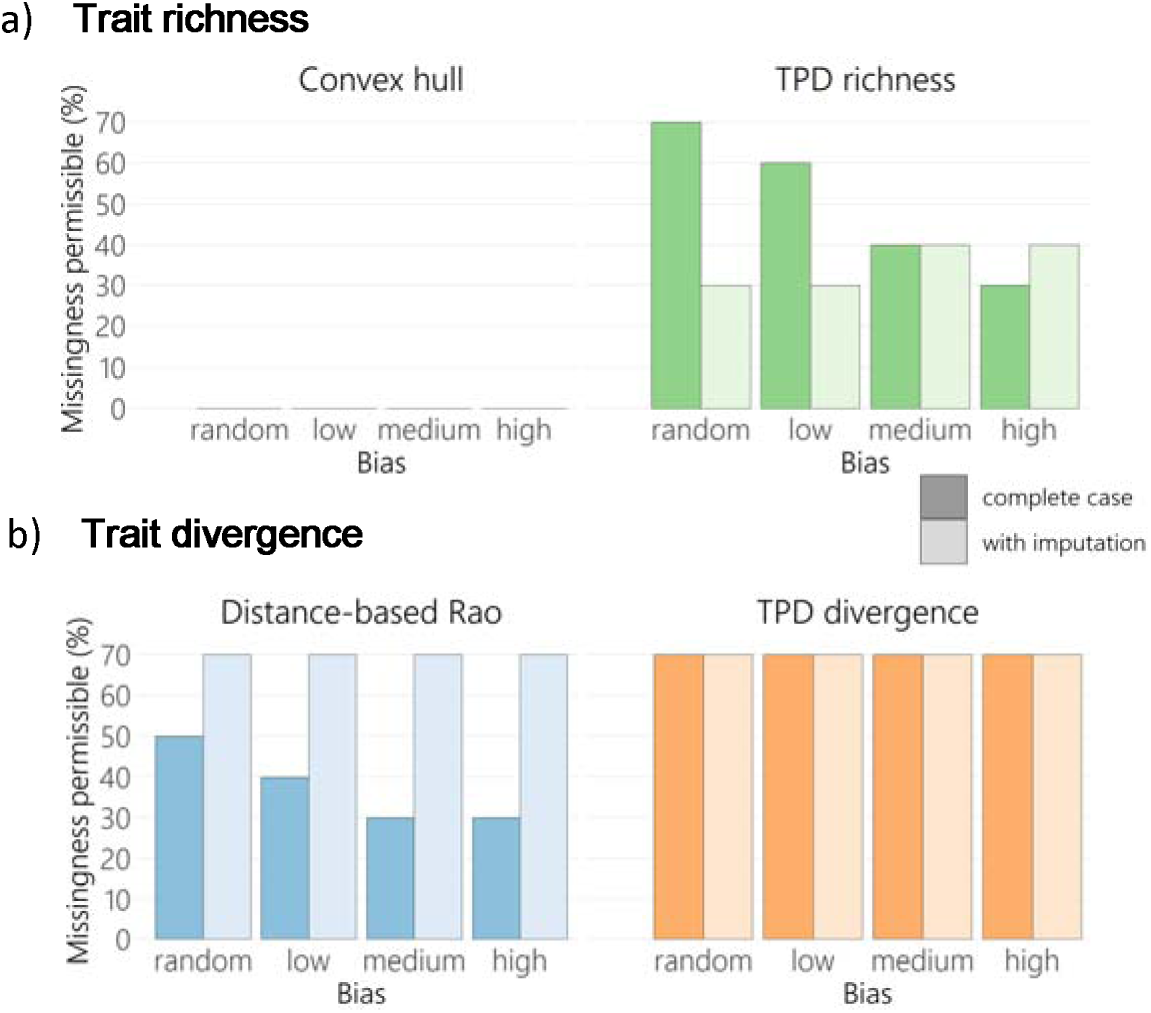
Maximum missingness permissible (maximum missingness where all trait diversity values fell within +/− 10% of trait diversity of the master dataset) for a) trait richness and b) trait divergence, with four degrees of bias. For each species with missing data, seven out of 11 traits were removed at random. Trait diversity was calculated using convex hull (richness), trait probability densities (TPD) richness, distance-based Rao (divergence) and trait probability densities (TPD) divergence. Results are shown for complete case analysis (darker colour bars) and for imputed datasets (lighter colour bars).

### 3. Quantifying bias in historic accumulation of large-scale bird trait datasets

We analysed three bird trait datasets to quantify the extent of bias and missingness present in real world datasets, with respect to geographic range size and body mass. These datasets were chosen because they all have 1000 species or more, and demonstrate the temporal accumulation of data, as Chira (2020) builds on Chira (2018), and Chira (2018) builds on Cooney (2017), using the same method of data collection. We plotted the probability density of the log_10_-transformed body mass and range size data and compared them to that of a full dataset (AVONET Version 3, Tobias et al., 2022). Body mass and range size data was obtained from AVONET Version 3 (Tobias et al., 2022).

The proportion of species included affected the probability distribution of body mass of species represented (Table S1). As data was accumulated (looking left to right in figure S5) the probability density distribution of included species appeared closer to that of the full dataset (blue in figure S5). Interestingly, bias affected the probability density of body mass of species present rather than truncating representation at extreme values: the smallest and largest body mass species were represented in every dataset, as shown by the points in figure S5. In contrast to included data, missing data became more skewed towards small species as the datasets became more complete (proportion of missing data decreased).

**Figure S5.**
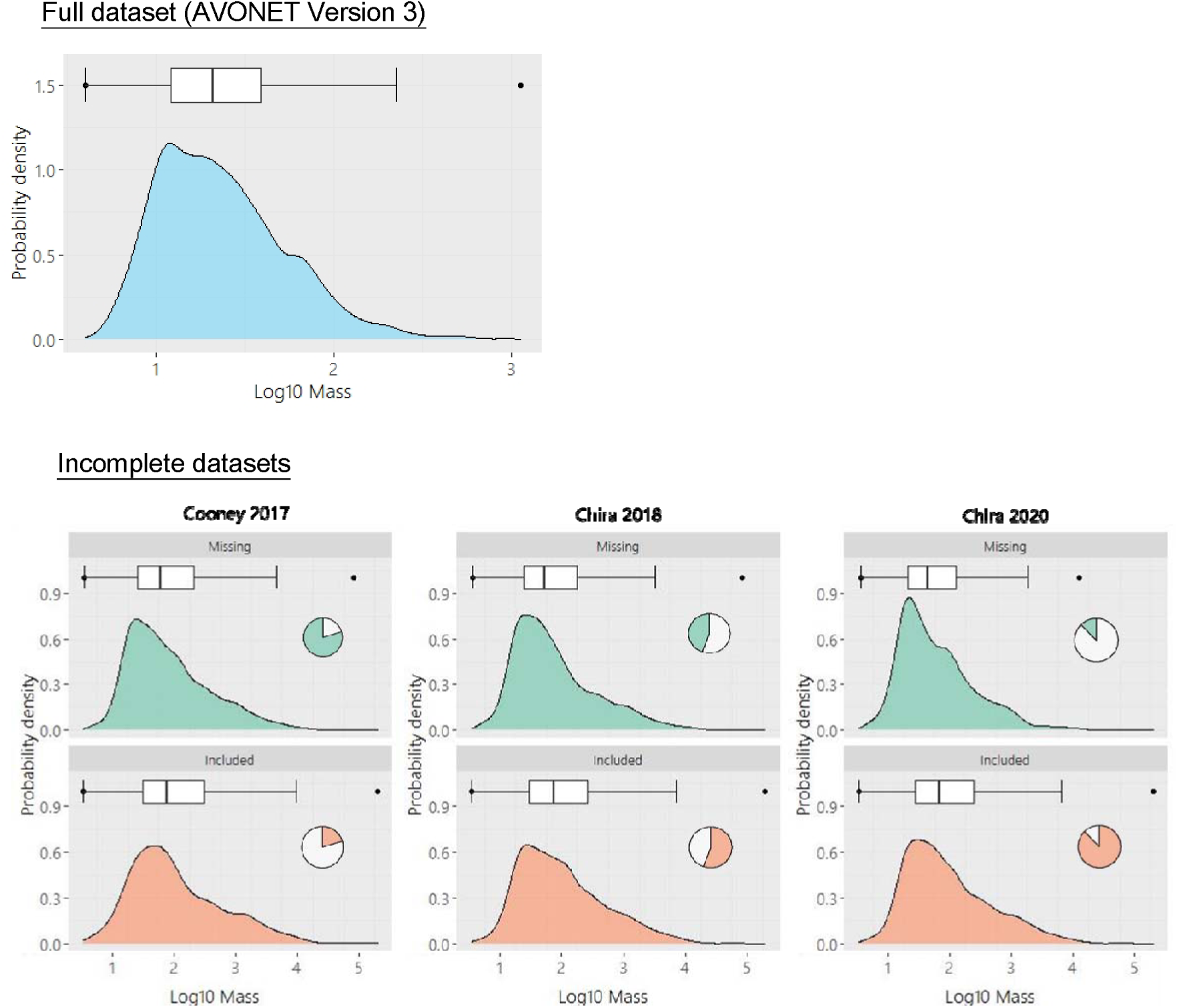
Probability density plot of body mass (log_10_ transformed) in three bird trait datasets for the full dataset (blue) and incomplete datasets, for species included (orange) and species missing (green). Pie charts show the number of species included (orange) or missing (green) in each study.

We quantified the extent of bias present in each dataset for body mass and range size (both log_10_ transformed, table S1, range size shown in figure S6). We did this by finding the probability that missing species had a body mass of lower than the median body mass of the full dataset (table S1). In this case a probability of 50% would represent an unbiased dataset, where species missing had an equal probability of occurring above and below the median of the full dataset.

**Figure S6.**
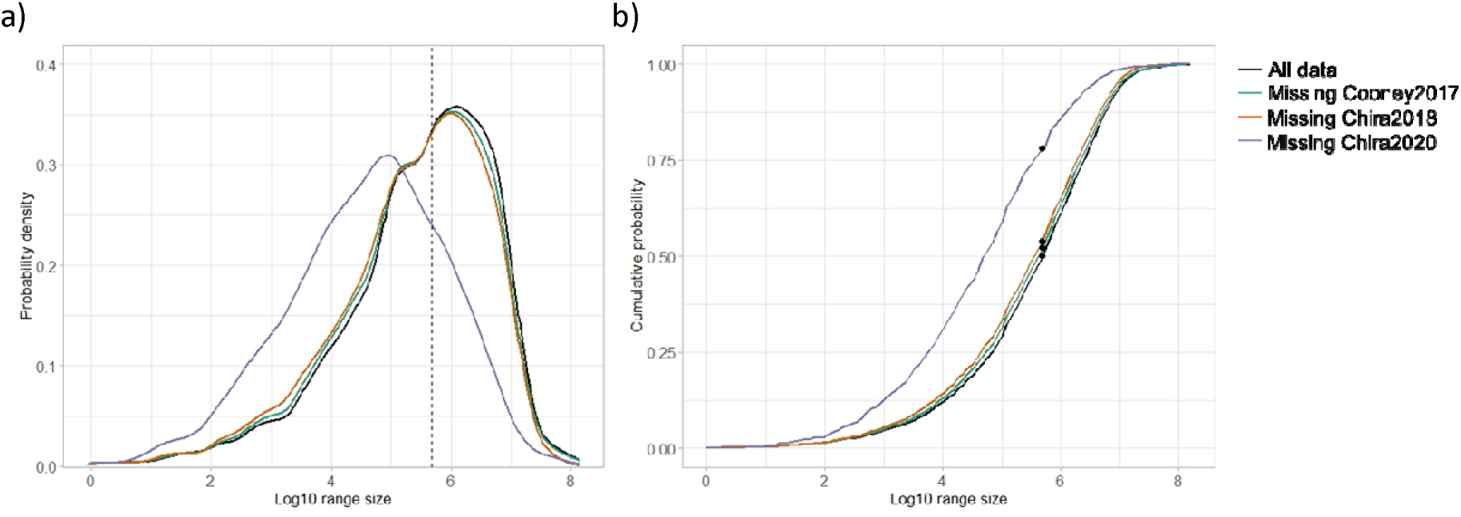
a) Empirical distribution function and b) cumulative distribution function of log10 transformed range size of species present in the full dataset (AVONET Version 3) and missing from the incomplete datasets (Cooney, 2017; Chira, 2018; Chira, 2020). The dotted line shows the median of the full dataset in a) and the points in b) show the probability of missing species occurring on or below the median of the full dataset.

**Table S1.** The probability that missing species have a body mass lower than the median body mass or range size of the full dataset (AVONET).

| Dataset | Species missing (%) | Probability missing data < median in full dataset (%) |  |
| --- | --- | --- | --- |
|  |  | Range Size | Mass |
| Cooney et al. (2017) | 80 | 52 | 51 |
| Chira et al. (2018) | 45 | 54 | 56 |
| Chira et al. (2020) | 13 | 78 | 60 |

From these results we chose to analyse scenarios of bias between 50% and 80% (probability that missing data were removed from species with below median body mass). As the maximum observed bias in the three datasets was 78% for range size, and 60% for body mass, 80% represents an extreme scenario of bias. Therefore, it is anticipated that the range of bias intensity present in real world datasets will be covered by these scenarios.

### 4. Error introduced by principal component analysis

To remove error introduced from conducting principal component analysis (PCA) independently on each dataset, we projected trait values from complete-case and imputed datasets onto the principal components obtained from the master dataset. To do this, for each principal component, we multiplied every trait value by the rotation given in the master PCA. For each species and principal component, trait values, multiplied by rotation, were summed to give the projected principal component value. For every dataset, we compared calculated (used in the main text) and projected principal components by estimating the correlation between the two trait spaces with a Procrustes test. We assessed the significance of the correlation using permutation tests (with 9999 randomizations) based on Monte Carlo simulations (using the *procuste.rtest* function from the *ade4* package [Dray & Dufour, 2007]). For all datasets we found high correspondence between calculated and projected principal components (p<0.0001 for all Procrustes tests).

We calculated trait richness for all datasets with calculated and projected principal components. For TPD, the NRMSE of trait richness estimated using calculated principal components compared to projected principal components was 4% of the range of trait richness values across all datasets. For convex hull NRMSE was 1% of the range in trait richness values across all datasets. Due to high correspondence between calculated and projected principal components, and low error in trait richness when using calculated principal components, we concluded that the response of PCA to incomplete datasets was limited and was not an important factor in determining trait diversity deviations.

### 5. The impact of tree balance on imputation accuracy

Biased removal of data was expected to increase tree imbalance, which given the use of phylogenetic information in missForest imputation, could affect imputation accuracy. To test this, we assessed how tree imbalance was affected by missingness for different scenarios of bias. We used two metrics to quantify tree imbalance; the corrected Colless index (Heard, 1992) and average leaf depth (Shao & Sokal, 1990). We chose these metrics because they are normalised by tip number so are independent of the number of species in the tree and they are based on the Colless (Shao & Sokal, 1990) and Sackin index (Shao & Sokal, 1990) respectively, which are widely used and accepted metrics (Fischer et al., 2021). We dropped tips for species with missing data, from a binary rooted maximum clade credibility tree, created using the first 1000 trees from Jetz et al. (2012).

When using the Colless corrected index, tree imbalance increased (figure S7) as more species were removed, but this effect was more subtle for high bias scenarios. This is likely because of the clustering of small species in trait space, so when species removal was biased the tree became less imbalanced than when species were randomly removed.

**Figure S7.**
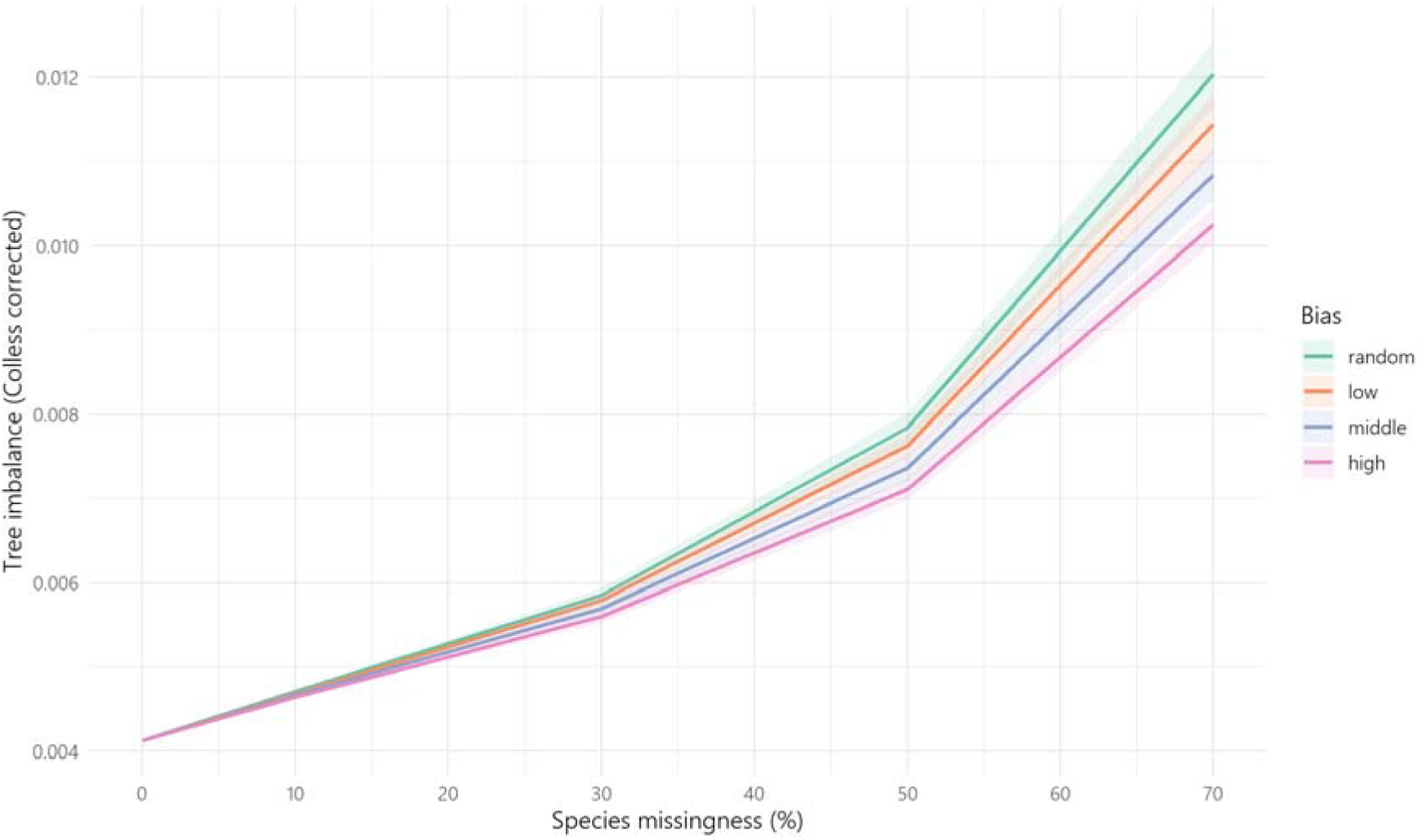
Tree imbalance (calculated with the Colless corrected index) with varying species missingness (% of species removed from tree) and bias (probability that species removed had a below median body mass).

In contrast, average leaf depth decreased in response to an increase in missingness and this response was greater in a high bias scenario (figure S8). In contrast, average leaf depth decreased in response to an increase in missingness and this response was greater in a high bias scenario (figure S8). The response to bias appeared to be similar as the corrected Colless index, but with a different direction of response to species removal at random.

**Figure S8.**
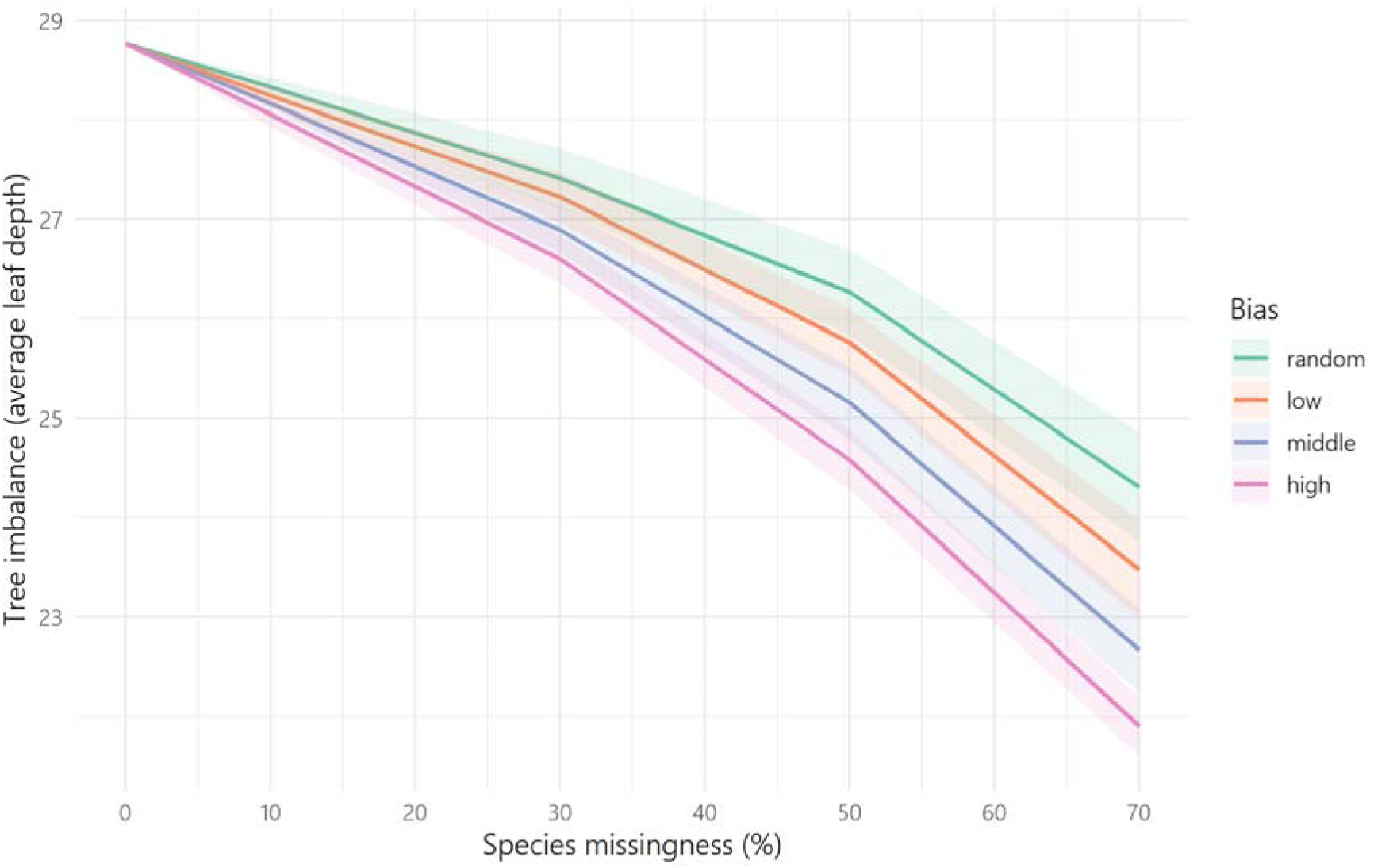
Tree imbalance (calculated with the Colless corrected index) with varying species missingness (% of species removed from tree) and bias (probability that species removed had a below median body mass).

We also tested whether tree imbalance affected imputation accuracy. Without grouping the results by missingness, there was little to no relationship between tree imbalance and imputation error (figure S9). However, when using the corrected Colless index and grouping by species missingness and bias there was a subtle positive relationship between tree imbalance and imputation error (**figure S10**). This does however appear to be a minor factor affecting imputation error relative to species clustering in trait space (resulting in high imputation accuracy for small species). The opposite relationship was observed when using average leaf depth, with a negative relationship between tree imbalance and imputation error (figure S10). It is possible that the limited impact of tree imbalance on imputation accuracy was due to the large number of species present in the tree, meaning that the range of tree imbalance values was very low even across missingness and bias scenarios.

**Figure S9.**
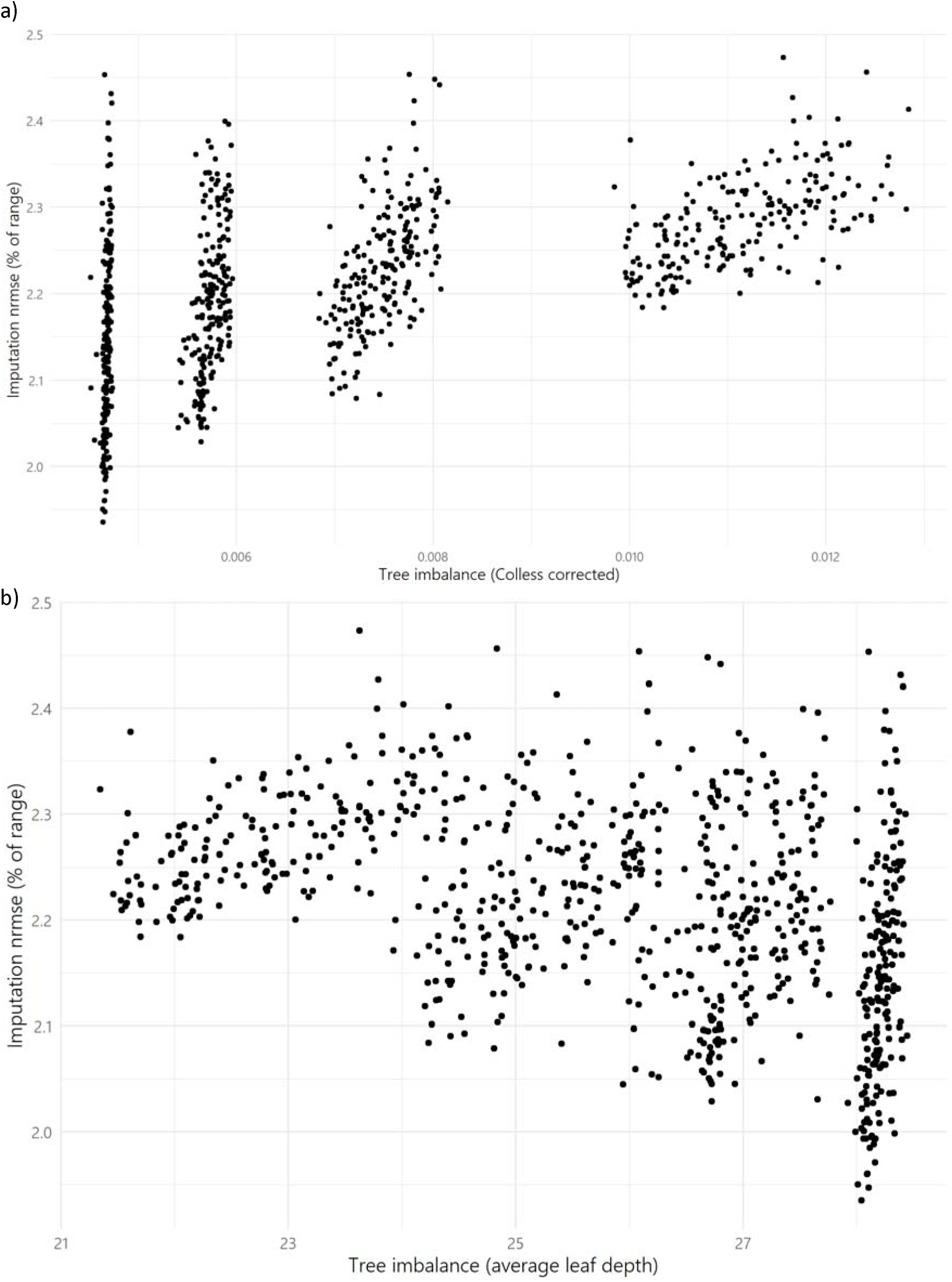
Tree imbalance calculated with a) Colless corrected index and b) average leaf depth against imputation accuracy (nrmse, % of range).

**Figure S10.**
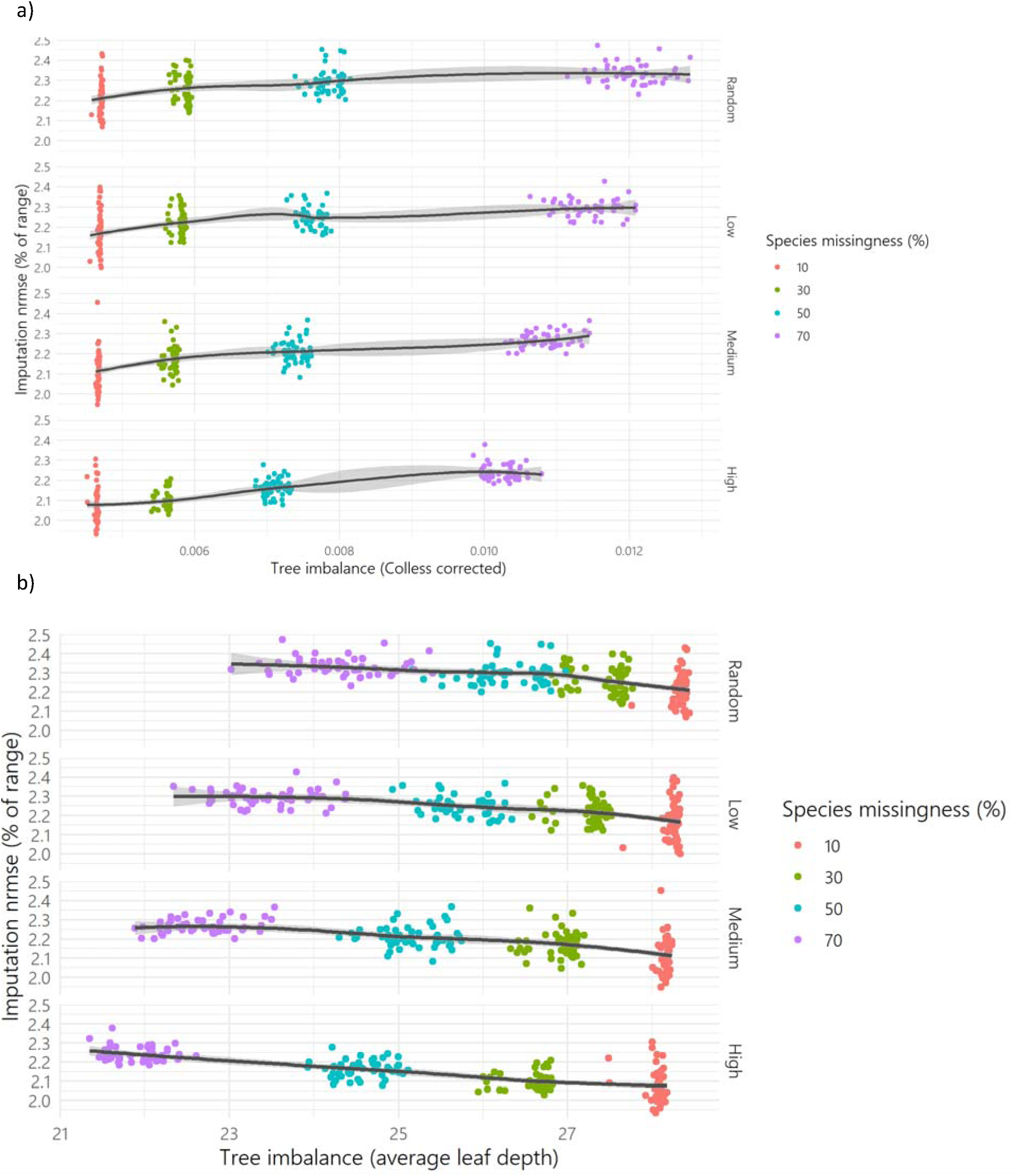
Tree imbalance calculated with a) Colless corrected index and b) average leaf depth against imputation error (nrmse, % of range), coloured by species missingness (% of species removed) for different scenarios of bias (random, low, medium and high). The black line shows a loess regression between tree imbalance and imputation error.

### 6. Increased trait richness under biased removal of species when using TPD

Trait richness when calculating according to TPD showed little response to random data removal but increased when data removal was biased towards small species. Intuitively data removal is expected to decrease trait richness. However, as species are removed, variance in the dataset increases, and smoothness of trait probability densities increase (as the bandwidth, the parameter determining smoothness in kernel density estimation, is selected accorded to the variance in the dataset). This can result in an increase in the trait range included in 95% of the probability density function (figure S11).

**Figure S11.**
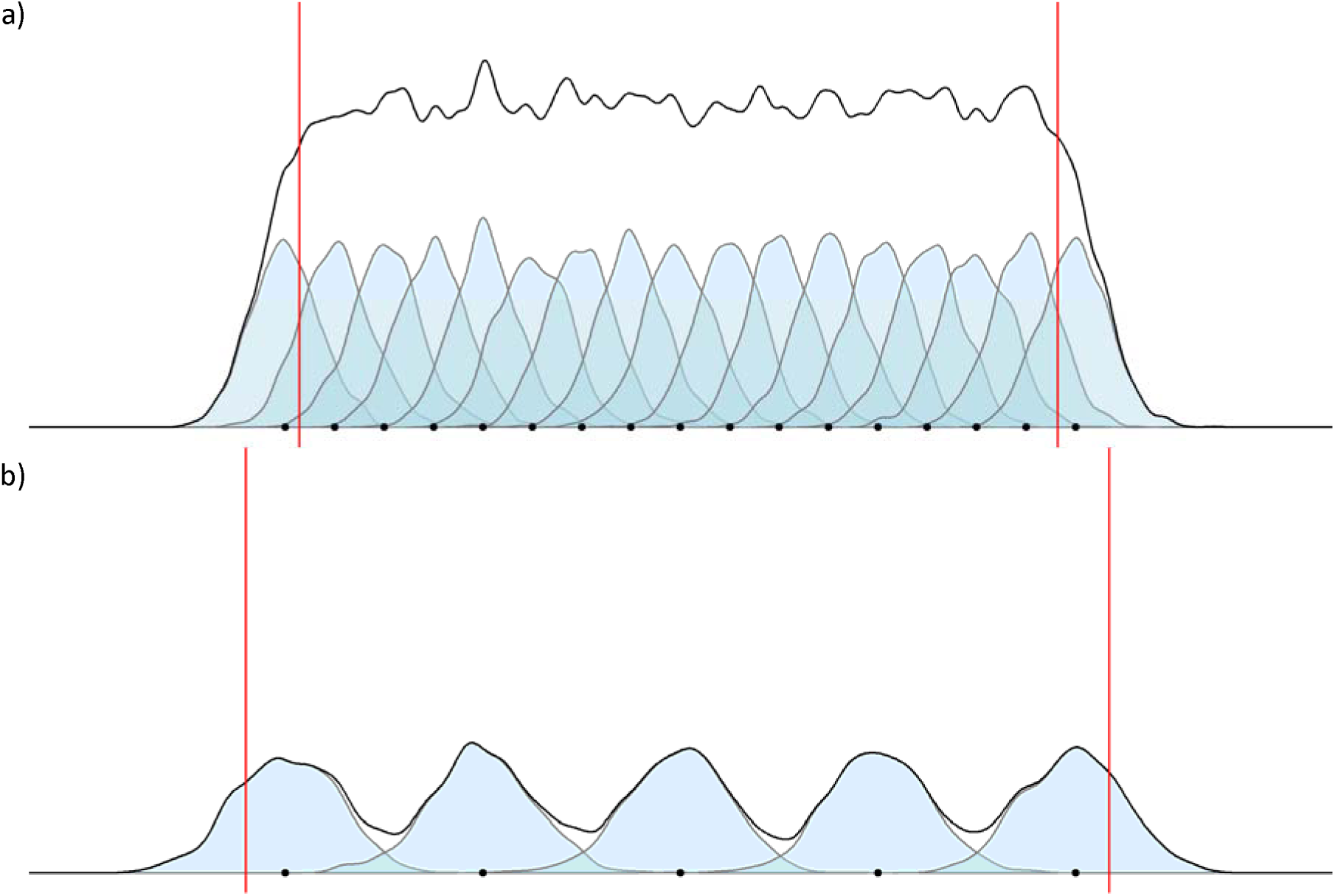
Conceptual diagram demonstrating how data removal (b) and increase in sample variance can result in an increase in trait range included in the 95^th^ percentile of probability density functions (red line) (relative to the master dataset, a). Dots hypothetical trait data along one trait axis, blue areas show the trait probability densities estimated for single data points, and the black line shows the convolution of these probability densities, to give the trait probability of the sample. Hypothetical trait data in a) have the same range as those in b).

This can be seen in real data (figure S12). In scenarios of high missingness and bias, 95% percentiles of probability densities calculated from incomplete datasets are often outside those of the complete dataset.

**Figure S12.**
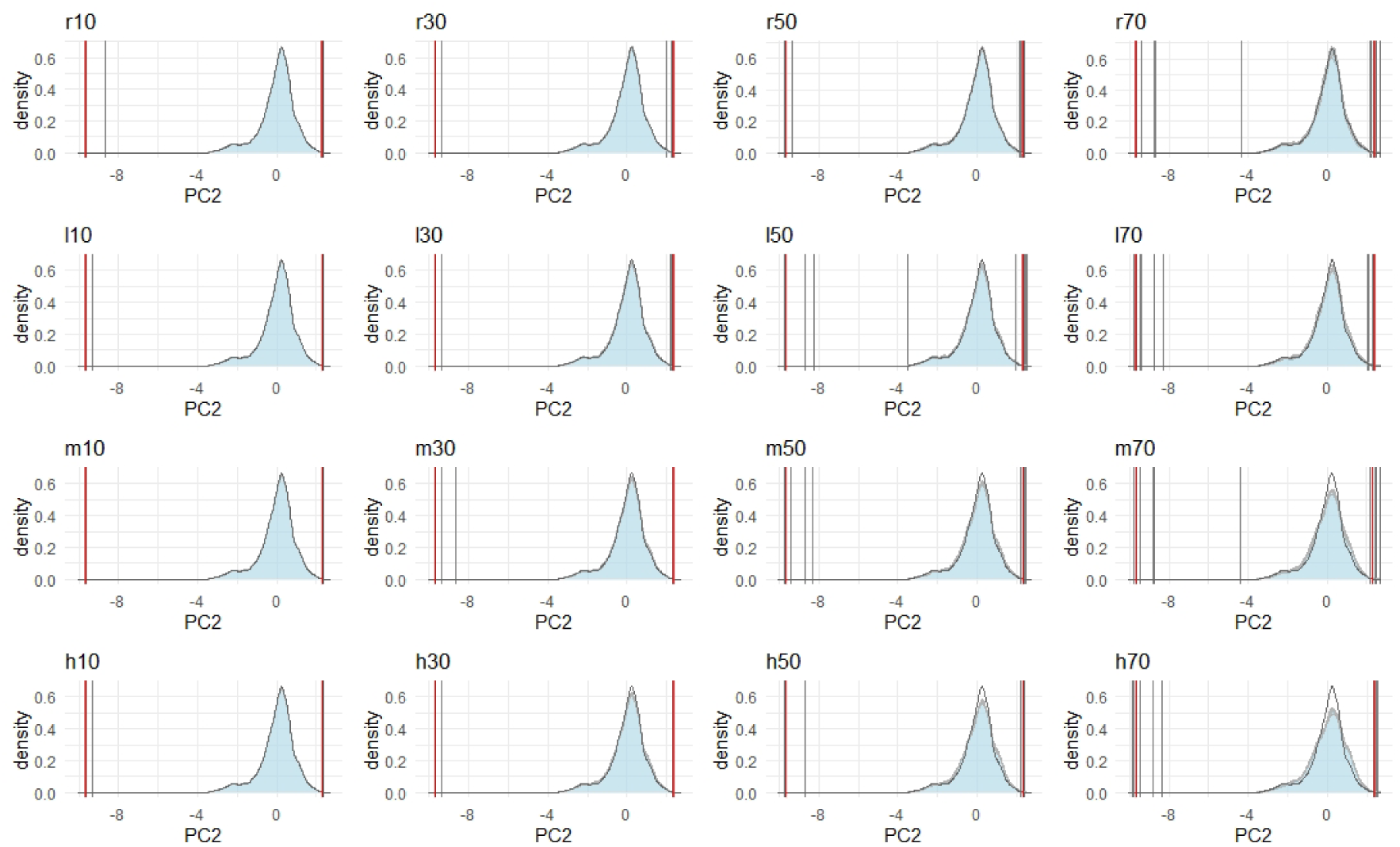
Probability distributions of principal component 2 (PC2) in incomplete datasets (blue area and grey lines. The black line shows the probability distribution of principal component 2 in the full dataset. Red lines show the 5^th^ and 95^th^ percentile of the full dataset, grey vertical lines show 5^th^ and 95^th^ percentiles of the probability density functions of incomplete datasets. Where grey lines are outside red lines, this shows that trait richness has increased as a result of data removal. Bias scenarios are indicated by letters above plots (r= random, l=low, m=medium, h=high) and missingness by the numbers (referring to % of species removed). 10 replicates for each scenario are shown.

This response can be suppressed by setting a constant bandwidth of kernel density estimation. Here we demonstrate this by setting bandwidth of the incomplete datasets to that of the full dataset. This results in a decrease in trait richness with data removal, as expected (figure S13).

**Figure S13.**
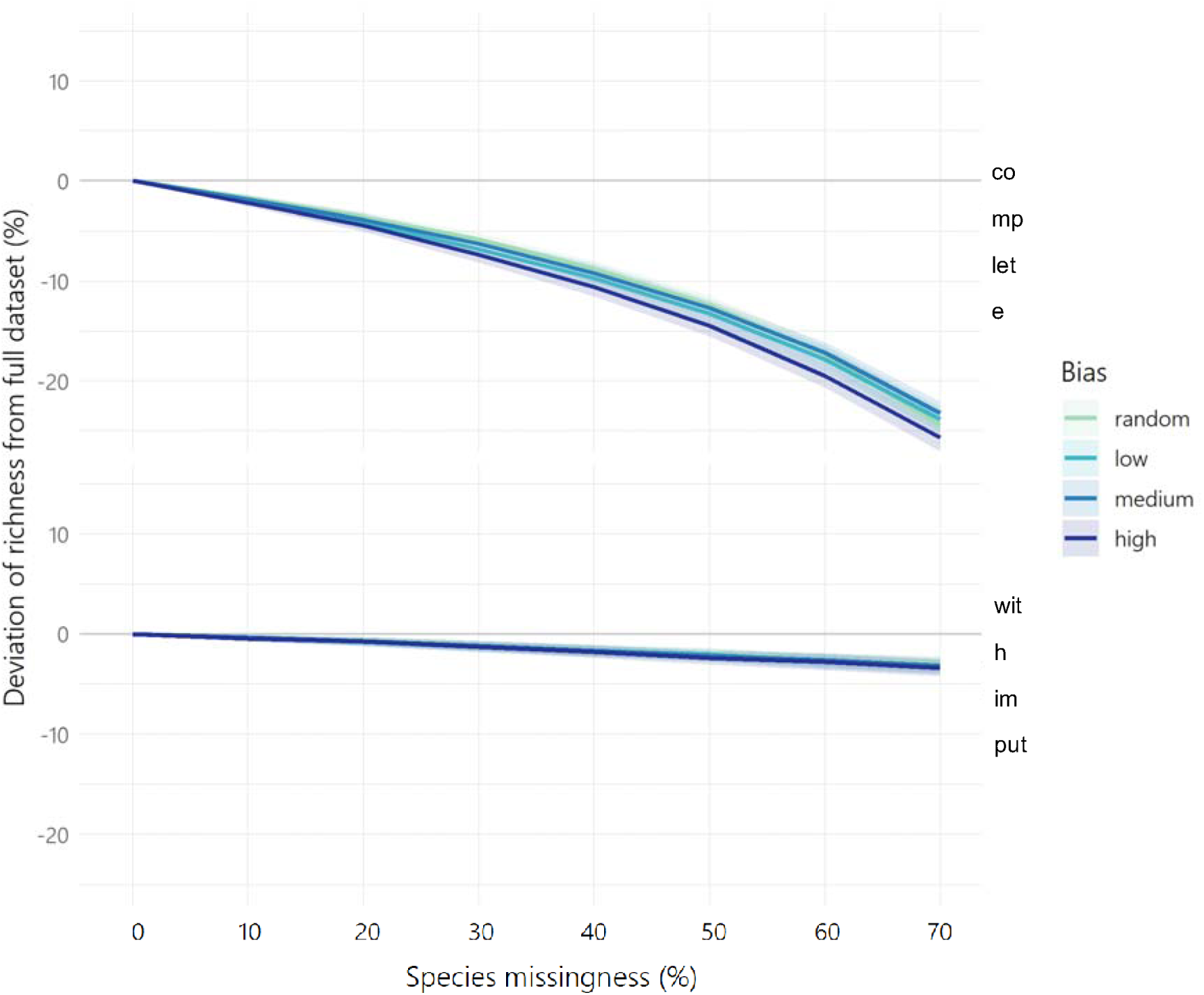
Deviation in trait richness of datasets with missing trait data compared to the complete dataset (mean shown by line with shaded areas showing 1 SD from 100 replicates) given varying levels of missingness (proportion of species with missing data) and bias (greater probability of data missing from small-sized species). The top panel shows results for complete case analyses (only species with all trait data analysed), and the bottom panel shows results for when missing values were imputed using Phylopars. Trait richness was calculated using trait probability densities (TPD) and bandwidth of kernel density estimation was fixed equal to the bandwidth estimated for the full dataset.

The bandwidth of kernel density estimation can be fixed when removal of species is simulated, for example to estimate the impact of extinction, but this is not sufficient to emulate the trait richness of a complete dataset given an incomplete dataset as the variance of the complete dataset will be unknown. As such trait richness of incomplete datasets may be overestimated by up to 10% when using TPD, depending on the extent of missingness and bias.

